# Ribosomal protein L5 (RPL5/uL18) I60V mutation is associated to increased translation and modulates drug sensitivity in T-cell acute lymphoblastic leukemia cells

**DOI:** 10.1101/2025.07.16.665036

**Authors:** Lorenza Bacci, Daniela Pollutri, Israt Jahan Ripa, Michael D’Andrea, Virginie Marchand, Yuri Motorin, Anne-Marie Hesse, Yohann Couté, Kamil Filipek, Marianna Penzo

## Abstract

Somatic mutations in ribosomal proteins (RPs), including RPL5, have been reported in approximately 10% of pediatric patients with T-cell acute lymphoblastic leukemia (T-ALL). In cancer, the incorporation of mutant RPs into ribosomes often disrupts canonical ribosome function, thereby contributing to disease development. In this study, we aimed to characterize the effects of the RPL5-I60V mutation in the context of T-ALL, focusing on its impact on translation and cellular responses to a panel of compounds in vitro. Using CRISPR-Cas9, we generated a homozygous knock-in mutant in Jurkat cells and investigated its effects on ribosome biogenesis. We observed both quantitative and qualitative alterations in the production of the large ribosomal subunit. Ribosomes containing the mutant RPL5 protein exhibited intrinsically increased protein synthesis activity, which correlated with enhanced cellular proliferation. We then evaluated the response of these mutant cells to a panel of compounds targeting protein synthesis at various levels—including an MNK1 inhibitor, metformin, silvestrol, homoharringtonine, anisomycin, resveratrol, and hygromycin B—as well as cytarabine, a chemotherapeutic agent commonly used in T-ALL treatment. Our results showed that the RPL5-I60V mutation confers increased sensitivity to most of these compounds, with the exception of hygromycin B.

This study advances our understanding of how oncoribosomes contribute to cancer pathogenesis and highlights the therapeutic potential of directly or indirectly targeting altered ribosomes, offering insights for the development of personalized treatment strategies.

**Highlights:**

- RPL5 mutations have been reported in T-ALL
- Ribosomes containing the RPL5 I60V mutation exhibit increased translational activity
- Increased proliferation is observed in cells harboring the RPL5-I60V mutation
- RPL5-I60V confers a specific drug sensitivity profile

**Statements and Declarations:** Competing interests: The authors declare that they have no conflict of interest.

## 1. Introduction

T-cell acute lymphoblastic leukemia (T-ALL), accounting for ∼15% of pediatric ALL cases, is a highly aggressive hematologic malignancy marked by clonal expansion of immature T-lineage hematopoietic precursors infiltrating the bone marrow and expressing aberrant T-cell markers [1]. T-ALL is a molecularly heterogeneous disease, with onset and progression linked to the accumulation of genetic alterations in oncogenes and tumor suppressors involved in essential cellular functions [1]. Frequently mutated genes include T cell-specific transcription factors such as NOTCH1 (mutated in >60% of cases) [1], as well as genes encoding epigenetic regulators (e.g., PHF6), tyrosine kinases, and signaling proteins (e.g., JAK/STAT pathway components) [2]. Recently, somatic mutations in ribosomal protein (RP) genes have been identified as novel drivers in pediatric T-ALL pathogenesis [3,4]. Somatic mutations in RPL5, RPL10 [4], RPL11 [5] and RPL22 [6] have been described with a cumulative frequency close to 10% in pediatric T-ALL patients. RPs, along with rRNAs, form the ribosome’s structural core, and their dysregulation or mutation is linked to various hematologic and solid tumors [7,8]. Heterozygous large deletions or point mutations in RPL5 were identified in several sporadic human tumors, including breast cancer, glioblastoma multiforme and melanoma [9–11]. In addition to being a component of the ribosome large subunit (60S), with important structural and functional activities [12], RPL5 has also multiple ribosome-independent functions, among which the regulation of ribosomal stress responses [13], of DNA repair mechanisms [14] and of transcription [15]. Consequently, genetic alterations in RPL5 can perturb multiple cellular processes, ultimately conferring distinct oncogenic properties to cancer cells. A principal outcome of RP mutations is the disruption of canonical ribosome function, resulting in the emergence of so-called “oncoribosomes” [7]. These aberrant ribosomes are proposed to exhibit functional specialization, potentially enabling the preferential translation of specific mRNA subsets in a context-dependent manner, thus contributing to the tumorigenic process [16,17]. This translational reprogramming represents an additional layer of gene expression regulation contributing to tumorigenesis [18].

Although, in the context of childhood T-ALL, currently available therapies, such as chemotherapy, lead to long-term remission for over 90% of patients [19], relapse remains a relevant scenario. The main cause of relapse is the emergence of resistance mutations in leukemia cells [20]. Since most of the tumors are characterized by hyperproliferation and an increase in the rate of protein synthesis to sustain cellular growth, a growing list of compounds targeting different steps of translation and/or ribosome biogenesis has been tested, and in some cases approved, for the clinical treatment of different tumor types [21,22].

The development of drugs specifically targeting *oncoribosomes* would represent a promising therapeutic strategy in the perspective of personalized medicine [23]. In this study, we investigated the impact of mutated RPL5 on cellular phenotype and on the response to different therapeutic compounds, using a T-ALL cellular model in which we knocked in a prototypic RPL5 mutation (I60V). Our findings demonstrate that a single RPL5 mutation induces distinct phenotypic traits in cells, accompanied by changes in drug sensitivity that are strongly influenced by the compound’s mechanism of action.

## 2. Materials and methods

### 2.1 Cellular model setup

Jurkat cells carrying the RPL5 c.178A>G (p.I60V) mutation were generated by CRISPR/Cas9 genome editing using the Alt-R™ CRISPR-Cas9 System (Integrated DNA Technologies). Briefly, ribonucleoprotein (RNP) complexes were generated by combining 160 pmol of equimolar crRNA (custom, Table 1):tracrRNA duplex with 80 pmol of Cas9 endonuclease. RNP complexes and 5 μg of single strand HDR donor template (ssODN, Table 1) were mixed with 10^6^ cells to a final volume of 100 μl in EP buffer (Optimem medium, Thermofisher) and electroporated by using Nepa21 electroporator system (Nepa Gene Co., Ltd) using the following settings for poring: pulse 175V, 5 ms length, 50 ms interval, and manufacturer’s recommended protocol for transfer pulse. 48 h after electroporation, cells were checked for mutation knock-in efficiency by genotyping assay using a droplet digital PCR approach (QX200 Bio-Rad Droplet Digital PCR System) with probes distinguishing the WT from the I60V allele (TaqMan® SNP Genotyping AssaysThermofisher). Homozygous I60V-RPL5 growing clones were isolated by single cell cloning and ddPCR genotyping screening. RPL5 mutational status of selected clones was confirmed by Sanger sequencing (for primers sequences see Table 1).

**Table 1.**
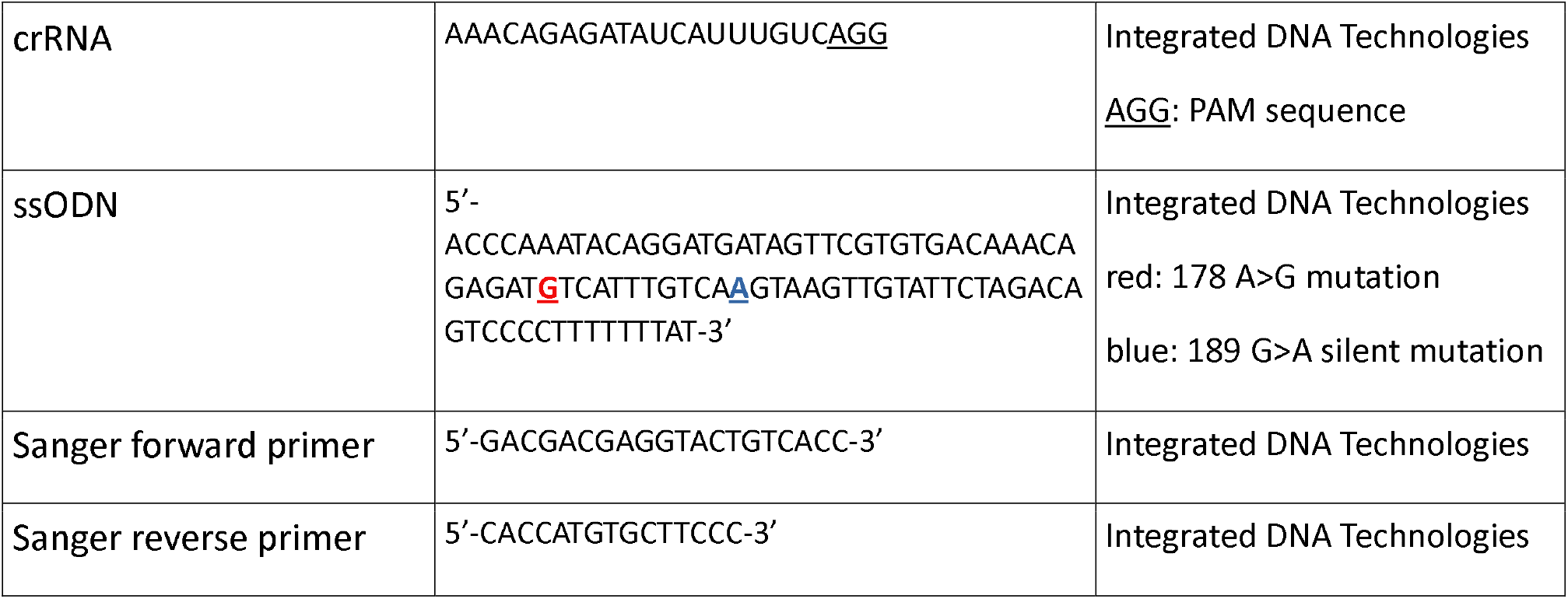
Custom synthesized sequences. *CrRNA: target-specific gRNA oligo, ssODN: knock-in single strand HDR donor template*.

### 2.2 Cell culture and growth curve

Jurkat clones were maintained in RPMI medium supplemented with 20% fetal bovine serum (FBS), 100 I.U./ml penicillin, 100 µg/ml streptomycin, and 2mM L-glutamine. Cells were grown at a maximum concentration of 10^6^ cells/ml. To overcome the biological variability intrinsic to clone-based experimental approaches, all experiments were carried out in parallel on 2 WT and 2 RPL5-I60V mutant clones.

For proliferation assay, a concentration of 3×10^6^ cells/ml was used for seeding the cells. Cell counting, performed in duplicate, was carried out every 24 hours by Countess 3 device (Invitrogen).

### 2.3 Polysome profiling

Cell suspension was treated with cycloheximide (CHX, Sigma-Aldrich) at a concentration of 100 μg/ml for 20 minutes at 37 °C and 5% CO_2_. The harvested cells were then washed with ice-cold PBS containing CHX (100 µg/ml, Sigma-Aldrich). Following this, the cells were lysed using a lysis buffer consisting of 0.1 M NaCl, 0.01 M Tris-HCl pH 7.6, 0.001 M EDTA, 0.1% Triton X-100, 0.1 mM PMSF, 100 µM Sodium orthovanadate, 100 mM DTT, and 100 µg/ml CHX, along with 1X Proteases inhibitor mix (Roche). The lysate was cleared at 12,000 g for 10 minutes at 4 °C. An amount of the cytoplasmic lysate containing 1 mg of total protein was loaded onto 15–40% sucrose gradients and centrifuged at 21,000 g at 4 °C for 2 hours using an Optima L70 ultracentrifuge (Beckman), with slow acceleration and deceleration settings. After centrifugation, the tubes containing the sucrose gradients were prepared for analysis at the Gradient Station (Isco Teledyne) and measured for absorbance at 254 nm. Simultaneously, the samples were divided into 2 ml tubes, with fractions collection every minute. For data analysis and visualization of the polysome profile, TRACERDAQ STRIPCHART software was employed.

### 2.4 RNA isolation from polysomal profile fractions and Bioanalyzer analysis

For each sample, the RNA preparation involved combining 1/10 of each fraction to generate a pool of cytoplasmic RNA. This was then treated with 100 μg/ml proteinase K (Sigma-Aldrich) and 10% SDS at 37 °C for 2 hours. Following digestion, a phenol: chloroform solution (5:1, pH 4.7) was added to each sample and mixed thoroughly. Additionally, 5 M NaCl was added and mixed to promote phase separation. The samples were then centrifuged at 16,000 RCF for 5 minutes at 4 °C. The upper aqueous layer was subsequently transferred to a fresh tube and 1 volume of isopropanol was added. The samples were thoroughly mixed and incubated overnight at -80 °C. Subsequently, centrifugation was performed at 16,000 RCF for 40 minutes at 4 °C, the supernatant was carefully removed, and the resulting pellet was washed with 70% ethanol. Finally, the dried pellet was resuspended in RNase-free water to prepare the nucleic acid samples for further analysis. 28S/18S ratio was calculated analyzing RNAs with a Bioanalyzer 2100 system (Agilent).

### 2.5 Ribosomes isolation

Ribosomes were purified as previously described [24]. Briefly, cells of two different WT and RPL5 I60V mutant clones were lysed in 10 mM Tris-HCl, pH 7.5, 10 mM NaCl, 3 mM MgCl_2_ and 0.5% (vol/vol) Nonidet P40 for 10 minutes on ice, and cytoplasmic lysates were collected upon centrifugation at 20,000 g for 10 minutes at 4 °C. After performing a 10 min incubation in protein synthesis master mix to achieve ribosomes run off from endogenous transcripts, lysates were loaded on a double cushion sucrose gradient with high salt/low salt conditions, to detach all ribosomes interactors. Ribosomes purification was achieved with a 15 h ultracentrifugation at 110000 g_av_ at 4 °C. Ribosomal pellets were resuspended in ribosome solution and ribosome concentration was quantified by measuring the absorbance at 260 nm (A_260_) in a NanoDrop®ND-1000 UV-Vis Spectrophotometer (Thermo Scientific) and applying the following formula: [mg/ml] of ribosomes = A_260_/12.5.

### 2.6 Cell free translation assay

This assay was carried out as described in [24], with minor modifications. The transcript used in these assays was a pR-CrPV_IRES-F transcript, previously transcribed *in vitro* as indicated in [25].

### 2.7 Mass spectrometry (MS)-based proteomic analysis of purified ribosomes

Proteomics analysis was carried out essentially as previously described [26] with minor modifications. In particular, peptide separation was performed on a 75 μm 250 mm C18 analytical column (Aurora Generation 2, 1.6 µm particles, IonOpticks). The chromatographic method consisted of a 120-minute multi-step gradient, increasing from 5% to 41% acetonitrile in 0.1% formic acid, at a flow rate of 400nL/min. Peptide and protein identification were performed with Mascot (v2.8), searching against the Uniprot human proteome database (March 2022 release), a custom sequence of the mutated RPL5 protein, an in-house contaminant database and their corresponding decoy versions for false discovery rate estimation. DDA results were merged using Proline software (v2.1). Validation and label-free quantification parameters followed those previously described [27]. Razor and specific peptides were used for quantification, except for RPL5 and its mutant form for which only strictly unique peptides were used. Mass spectrometry proteomics data have been deposited in the ProteomeXchange Consortium via the PRIDE repository under dataset identifier PXD064727 [28]. Statistical analysis was carried out using ProStaR 1.38.0 [29] following previously described method [30] with minor modifications, including an absolute log2 (fold change) threshold greater than 1.5 and an adjusted p-value (Benjamini-Hochberg correction) below 1%.

### 2.8 RiboMethSeq and HydraPsiSeq analysis of rRNA modification profiles

rRNA was extracted from purified ribosomes with PureZOL^™^ (BioRad) following manufacturer’s protocol. Prior to proceeding further, rRNA quality was assessed with a Bioanalyzer RNA 6000 Nano assay (Agilent): only samples with a RIN > 9 were selected for high throughput modification analyses. RiboMethSeq and HydraPsiSeq protocols used for quantification of rRNA modifications have been extensively described previously [31–36]. Both methods quantitatively measure protection of the modification-adjacent phosphodiester bond against chemically induced cleavage, which is alkaline hydrolysis for RiboMethSeq and hydrazine/aniline treatment for HydraPsiSeq. Analysis is performed simultaneously for all known modification sites (110 Nm and 107 ψ). In brief, rRNA is subjected to chemical treatment allowing to reveal resistant RNA phosphodiester bonds and the resulting fragments are converted to sequencing library using NEBNext Small RNA kit. Sequencing is performed in a single read SR50 mode, with the target value of ∼10-15 mln of raw reads for RiboMethSeq and 20-25 mln for HydraPsiSeq. Bioinformatics treatment pipeline consists in trimming of the adapter sequence, alignment of the trimmed reads to the reference sequence and counting of 5’/3’-end for RiboMethSeq and 5’-ends only for HydraPsiSeq. Combined end-coverage profile is further used for calculation of MethScore and PsiScore, respectively.

### 2.9 Compounds preparation

For each compound, the optimal concentration range, to be tested on Jurkat cellular model, was determined based on available literature data. All the compounds (mentioned in Table 2) were purchased from Merck Life Science. Compounds solutions were prepared by dissolving them in specific solvents (water, ethanol, DMSO, as indicated in Table 2) to prepare concentrated stock solutions. These stocks were further diluted freshly for each experiment.

**Table 2.** List of tested compounds. For each drug, the solvent used, along with the concentrations used in this study and relevant references, are indicated.

| Compound | Vehicle | Higher Dose | Intermediate Dose | Lower Dose | References |
| --- | --- | --- | --- | --- | --- |
| Homoharringtonine | DMSO | 1 μM | 0,1 μM | 10 nM | [37] |
| Cytarabine | Water | 1 μM | 0,1 μM | 0,01 μM | [38] |
| MNK1 Inhibitor<br>4-Amino-5-(4-fluoroanilino)-pyrazolo[3,4-d]pyrimidine | Ethanol | 100 μM | 10 μM | 1 μM | [39] |
| Resveratrol | DMSO | 500 μM | 100 μM | 10 μM | [40] |
| Anisomycin | DMSO | 10 μM | 1 μM | 0.1 μM | [41] |
| Metformin | Water | 10mM | - | 1 mM | [42] |
| Hygromycin B | Water | 750 μM | - | 500 μM | - |
| Silvestrol | DMSO | 1 nM | - | 0.1 nM | - |

### 2.10 AlamarBlue^™^ assay

For each clone, 1 × 10^5^ cells were seeded in a 96-well plate, in the presence of the drugs as indicated in Table 2, or in the presence of the solvent with the same final dilution as in the drug-treated wells (as a negative control). The plate was incubated at 37 °C, 5% CO_2_, and controlled humidity for 48 hours. Alamarblue^™^ reagent (ThermoFisher Scientific) was added directly into the culture media at a final concentration of 10%, and the plate was incubated for 4 hours at 37 °C, 5% CO_2_, after which fluorescence was measured in a Spark plate reader (TECAN).

### 2.11 Cell Viability Test

For each clone, 10^5^ cells were seeded in a 96-well plate, in the presence of the drugs at the following concentration: the MNK1 inhibitor at 1⍰µM, metformin at 1mM, silvestrol at 0.1⍰nM, homoharringtonine at 10⍰nM, anisomycin and cytarabine at 0.1⍰µM, resveratrol at 10⍰µM, and hygromycin B at 500⍰µM. Negative control were established using the solvent with the same final dilution as in the drug-treated wells. The plate was incubated at 37 °C, 5% CO_2_, and controlled humidity for 48 hours. RealTime-Glo™ kit solution (containing NanoLuc^®^ Luciferase and MT Cell Viability Substrate, Promega) was added at each well, following manufacturer’s specification. The plate was then incubated at 37 °C with 5% CO_2_ for 72 hours, and luminescence was recorded at specific time intervals in the Spark plate reader (TECAN).

### 2.12 Cytotoxicity and apoptosis

For each clone, 20,000 cells were seeded in a 96-well cell culture plate and treated with drugs at concentration reported in the previous paragraph and negative controls were established by seeding cells without drug treatment. The plate was then incubated at 37 °C, 5% CO_2_, and controlled humidity for 8 hours or 24 hours for apoptosis assessment, and 40 hours for cytotoxicity analysis. Then, the ApoTox-Glo™ Triplex Assay (Promega) was used to assay cytotoxicity or apoptosis, following the manufacturer’s instructions. Briefly, following incubation, a Viability/Cytotoxicity Reagent or Caspase-Glo^®^ 3/7 Reagent were added to assess cytotoxicity or apoptosis, respectively. Plates were incubated for 30 minutes at room temperature, then fluorescence or luminescence measurements were taken using a Spark plate reader (TECAN). For fluorescence measurements, the wavelength sets were 485 nm excitation and 520 nm emission for cytotoxicity. For luminescence measurement, integration time was set to 1 second.

### 2.13 SUnSET assay and western blot analysis

To assess protein synthesis, the SUnSET assay was carried out as previously outlined [43]. For each clone, 10^5^ cells were seeded in a 12-well cell culture plate and treated with drugs at the concentration reported in the previous paragraphs and negative controls were established by seeding cells without drug treatment. The plate was then placed in an incubator set at 37 °C, 5% CO_2_, and controlled humidity for 24 hours. Cells were exposed to puromycin (1 μg/ml) for 10 minutes (“pulse”) followed by two washes with ice-cold PBS. Subsequently, cells underwent a 50-minute “chase” period in fresh media at 37 °C with 5% CO_2_. Total proteins were extracted utilizing RIPA buffer (50 mM Tris-HCl pH 7.5, 150 mM NaCl, 1% IGEPAL, 0.1% SDS and protease inhibitor cocktail), and 30 μg of proteins were denatured and separated on a 10% polyacrylamide gel, then transferred onto nitrocellulose membrane, and protein loading was assessed through Ponceau staining. The membrane was then incubated overnight with the primary antibody [mouse anti-puromycin (12D10, Sigma-Aldrich), diluted 1:5000]. After incubation with HRP-conjugated anti-mouse IgG antibody (Jackson Immunoresearch) diluted 1:10000, the membrane blots were developed using the ChemiDoc Imaging System (Bio-Rad) and analyzed with Image Lab software.

### 2.14 Statistical analysis

Statistical analysis was performed using GraphPad Prism 8 statistical software. Statistical tests used and p-values are indicated in figure legends.

## 3. Results

### 3.1 Impact of I60V mutation on ribosome biogenesis and function

We selected I60V as a representative T-ALL–associated RPL5 mutation, since this substitution was previously identified in another T-ALL cell line derived from a pediatric patient (DND41) [4]. Indeed, I60 is located within the same functional region as other patient-derived mutations (Fig. 1A). RPL5 is assembled within nascent ribosomes in complex with RPL11 and 5S rRNA, in a structure called 5S ribonucleoprotein (5S-RNP). Even though the precise timing of incorporation of this complex in the large subunit during ribosome biogenesis is not clearly defined, it is known that it is a quite early event, occurring in the nucleolus (reviewed in [44]). On this premise, we reasoned that RPL5 I60V mutation could, in line of principle, impair the biogenesis of the large subunit, in terms of either RP or rRNA composition (Fig. 1A). To explore this specific issue, we generated a homozygous knock-in mutant model in Jurkat T-ALL cells and verified the genetic status by Sanger sequencing (data not shown). We then adopted multiple approaches to study ribosome biogenesis. First, we confirmed by MS-based quantitative proteomic analysis the incorporation of the I60V RPL5 protein in cytoplasmic ribosomes of mutant cells (Fig. 1B). This approach demonstrated that RPL5 was the only ribosomal protein differentially abundant in ribosomal fractions from WT and mutant cells, with WT RPL5 significantly enriched in WT cells and I60V RPL5 significantly enriched in mutant cells (Fig. 1B). In addition, few non-ribosomal proteins were found differentially abundant in ribosomal preparations from WT and mutant cells (Table 3). Polysome profiling indicated a slight decrease in the 60S and 80S peaks in RPL5-I60V compared to RPL5-wt (Fig. 1C, D), mirrored by a non-significant reduction in 60S/40S ratio (Fig. 1E), suggesting a quantitative effect of the RPL5 mutation on large subunit biogenesis. Consistent with this observation, we could report a decrease in the 28S/18S rRNA ratio determined on the rRNAs purified from total cytoplasmic RNA (Fig. 1F). Extensive rRNA modifications play an important role in the processing of rRNA itself, *2*’-*O*-Methylation and pseudouridylation being the most frequent modifications on rRNAs [45]. We therefore performed an analysis of rRNA *2*’-*O*-methylation and pseudouridylation profiles by RiboMethSeq [31] and HydraPsiSeq [32] protocols, respectively. Both methods have been fully optimized for quantitative assessment of the RNA modification level [33–36] and extensively used for analysis of rRNA modification dynamics. The results (Fig. 1G, H) clearly indicate that the I60V RPL5 mutation does not affect either modification.

**Table 3.** List of ribosomal and non-ribosomal proteins differentially expressed in wild type (WT) or mutant (Mut) clones by MS-based quantitative proteomic analysis. In red background, proteins that are up-regulated in I60V-RPL5 mutant clones, in blue background the down regulated.

| Protein Entry Name | Gene Name | Log<sub>2</sub> Fold Change (Mut vs WT) | p-value |
| --- | --- | --- | --- |
| RL5_mutI60V | RPL5mut | 8,19 | 1,9E-06 |
| Q16577_HUMAN | PTI-1 | 5,60 | 6,2E-04 |
| ARL4C_HUMAN | ARL4C | 3,44 | 5,8E-06 |
| XPF_HUMAN | ERCC4 | 3,38 | 4,6E-06 |
| H7C3W8_HUMAN | LRPPRC | 2,80 | 1,6E-04 |
| RL5_HUMAN | RPL5 | -6,46 | 3,8E-06 |
| NPC1_HUMAN | NPC1 | -2,87 | 2,9E-05 |
| ABCBA_HUMAN | ABCB10 | -2,57 | 8,7E-05 |
| ETFD_HUMAN | ETFDH | -1,71 | 3,3E-04 |
| B3GN2_HUMAN | B3GNT2 | -1,70 | 5,5E-04 |
| TRI27_HUMAN | TRIM27 | -1,53 | 1,2E-03 |

**Fig. 1.**
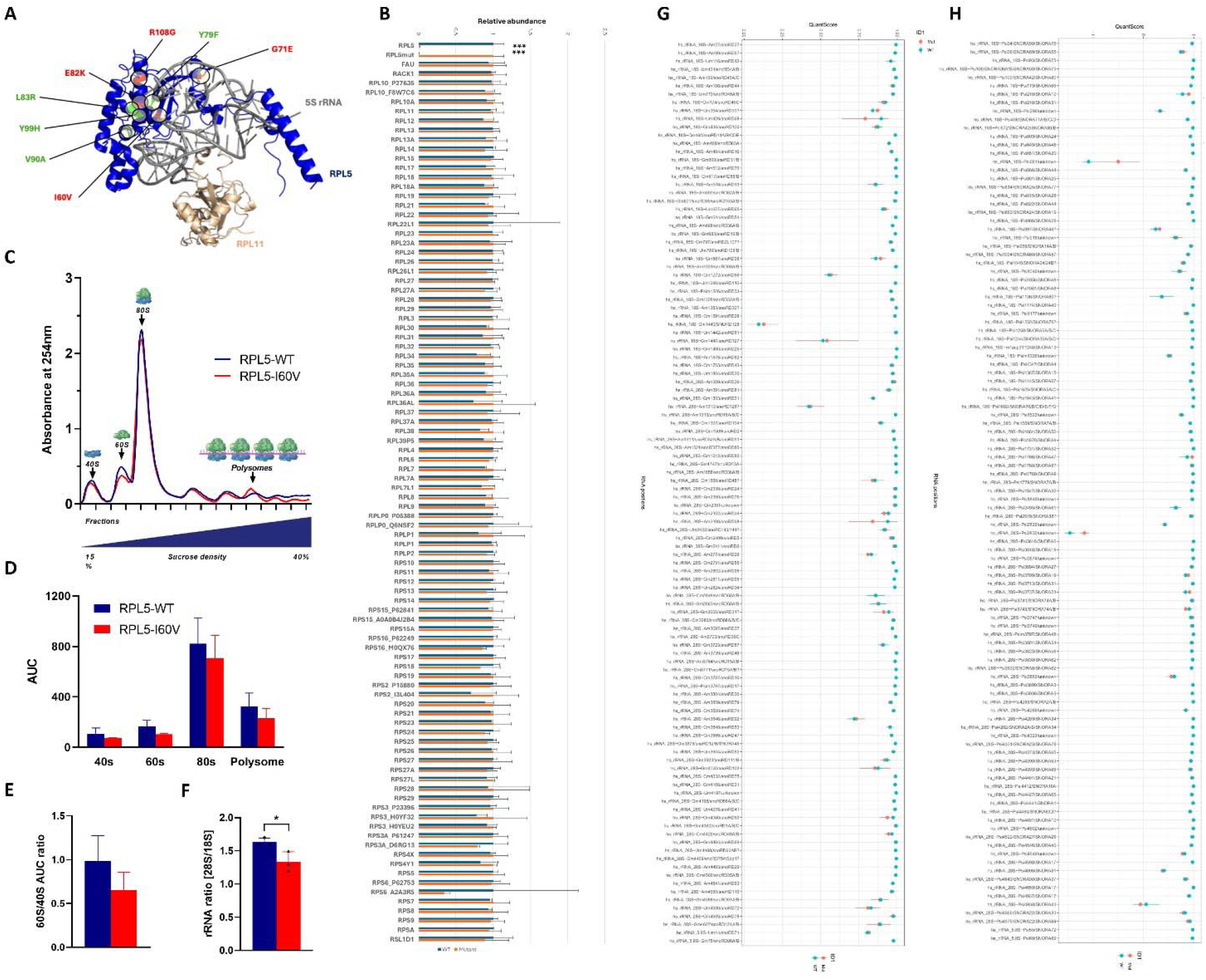
Impact of RPL5 I60V mutation on ribosome biogenesis. (A) RPL5 structure within the 5S-RNP. RPL5 (in blue) assembled with RPL11 (in gold) and part of the 5S rRNA (in grey), from the PDB structure 4v6x. I60 as well as the residues found mutated in pediatric T-ALL (from https://www.stjude.cloud/) are highlighted in red, whereas the residues found mutated in other pediatric cancers are in green. (B) Relative abundance of the RPs detected in MS-based proteomic analysis of purified ribosomes from 2 wild type clones (WT) and 2 I60V mutant (Mutant) clones. Data represent the mean of three independent experiments; for each protein, the maximum mean abundance was set to 1. Error bars represent standard deviation. ***limma p-value < 0.001 in the statistical analysis of quantitative proteomics data (see Materials and methods section). (C) Representative polysome profiles obtained by the separation on 15-40% sucrose-density gradients of cytoplasmic extracts from 2 WT and 2 RPL5 I60V clones. The average A_254_ values for each WT and mutant clone are shown in blue and red, respectively. (D) For each of the different samples, areas under the curve (AUC) were determined and plotted. (E) Ratio of 60S and 40S peak areas/total areas. (F) Ratio of 28S/18S total cytoplasmic rRNA, determined by Bioanalyzer. (G) RiboMethSeq analysis. Panel shows values of rRNA MethScore (QuantScore) for WT (blue) and RPL5 I60V cells (Mut, red). Analysis was performed for 2 WT and 2 mutant clones. The mean value is shown as a dot and error bar shows variability between two replicates. The majority of Nm sites are almost fully modified, while few show only partial methylation (MethScore <0.75). Identity of the rRNA Nm site as well as associated C/D-box snoRNA are shown at the bottom. (H) HydraPsiSeq analysis. Panel shows values of rRNA PsiScore (QuantScore) for WT (blue) and RPL5 I60V cells (Mut, red). Analysis was performed for 2 WT and 2 mutant clones. The mean value is shown as a dot and error bar shows variability between two replicates. Three rRNA sites with negative PsiScore levels show higher cleavage compared to the neighboring nucleotides and thus are not modified even in the WT human cells. Identity of the rRNA Nm site as well as associated H/ACA-box snoRNA are shown at the bottom.

We next evaluated the effect of the mutation on ribosomal function. Mutant cells showed an increase in puromycin incorporation in nascent peptides (Fig. 2A), indicating a more active protein synthesis. Consistently, we found that ribosomes purified from mutant cells had an intrinsic activity boosted by 50% more, compared to WT (Fig. 2B), and this higher synthetic activity proved to be independent of the initiation mode (cap vs IRES). In keeping with a more active translation, we found that RPL5 mutant cells had a higher growth rate compared to WT (Fig. 2C).

**Fig. 2.**
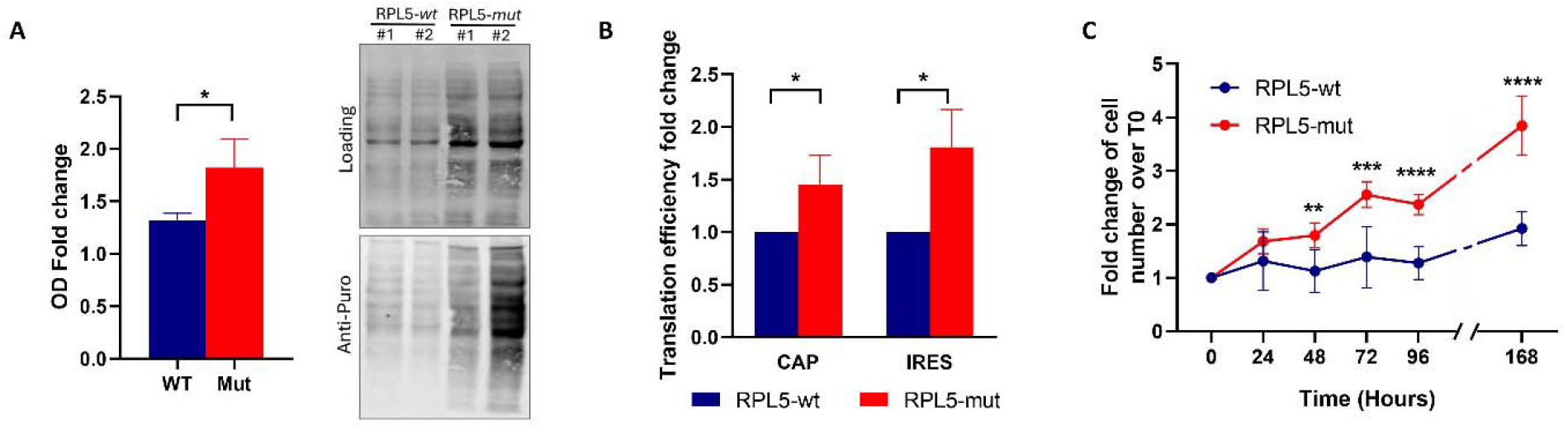
Impact of RPL5 I60V mutation on ribosome function. (A) Cellular protein synthesis rate tested by SUnSET puromycin incorporation. Right: representative western blot. Left: normalized densitometric quantification of puromycin incorporation. Data are plotted as fold change vs WT ± SD. (B) Cell free translation efficiency of ribosomes purified from WT or I60V mutant clones, quantified as cap-or CrPV IRES-mediated translation of a bicistronic Luciferase reporter mRNA. (C) Average proliferation curves over a 7 days-time span of WT and I60V mutant clones. Unless otherwise stated, experiments were performed in 3 biological replicates with 2 I60V and 2 WT clones. Statistical significance was calculated by t-test, with *p<0.05, **p<0.01, ***p<0.001 and ****p<0.0001.

### 3.2 Impact of I60V mutation on the efficacy of compounds with diverse mechanisms of action

Since our data indicated that the I60V mutation in RPL5 can significantly impact ribosome biogenesis, translation and, therefore, cellular proliferation, we next aimed to understand if the mutation could affect the effectiveness of drugs that target the translational machinery. Our interest was specifically focused on addressing whether the RPL5 mutation could, directly or indirectly, modulate the sensitivity of the cells to treatments. We therefore selected several drugs with different mechanisms of action on translation (ribosome biogenesis inhibitors, inhibitors of pathways upstream of protein synthesis, direct inhibitors of ribosome function). In addition, we added compounds used for T-ALL therapy, like cytarabine [46]. We tested the effect of these drugs on cellular metabolic activity, viability, protein synthesis and cell death in the WT or RPL5-I60V background. To assess their efficacy, we undertook an extensive literature review aimed at pinpointing optimal concentrations, in terms of cell growth inhibition, for our cellular model. These concentrations were selected as intermediate values within a wider spectrum, ranging from tenfold higher to tenfold lower concentrations (see Table 2). Since precise concentrations of hygromycin B and silvestrol were not readily available in the literature, a preliminary assessment was conducted to determine their effective concentration range (data not shown).

#### 3.2.1 Effect of the mutation on drug-mediated modulation of cellular metabolism

We first assessed the metabolic differences at the steady state for the cells harboring the two genotypes, finding a slight, non-statistically significant reduction in the I60V genotype (data not shown). We then defined the optimal drug concentration by evaluating the effect of drug treatments over 48 hours on cellular metabolic activity and viability by AlamarBlue^™^ assay. Based on their effect, compounds can be divided into two groups (Fig. 3): one for which there is a dose/response effect (MNK1 inhibitor, metformin, silvestrol, anisomycin, cytarabine) and one for which the effect is very similar at the different doses tested (homoharringtonine, resveratrol, hygromycin B). This grouping is independent of the genetic background (WT or I60V) of the cells (Fig. 3). As illustrated in Fig. 3, RPL5-I60V clones generally demonstrated equal or higher sensitivity to the different treatments, compared to WT except for hygromycin B, for which RPL5-I60V clones exhibited resistance at the doses tested. Based on these results, we could define, in the tested range, the optimal concentration to be employed for further experiments, i.e., the lowest possible concentration at which significant differences in metabolism were observed between the mutant and wild-type (WT) cells. Specifically, the observed differences occurred at 1⍰µM for the MNK1 inhibitor, 1⍰mM for metformin, 0.1⍰nM for silvestrol, 10⍰nM for homoharringtonine, 0.1⍰µM for anisomycin, 10⍰µM for resveratrol, and 500⍰µM for hygromycin B. In contrast, for cytarabine, the most notable metabolic decrease occurred at the intermediate concentration of 0.1⍰µM, which was therefore chosen.

**Fig. 3.**
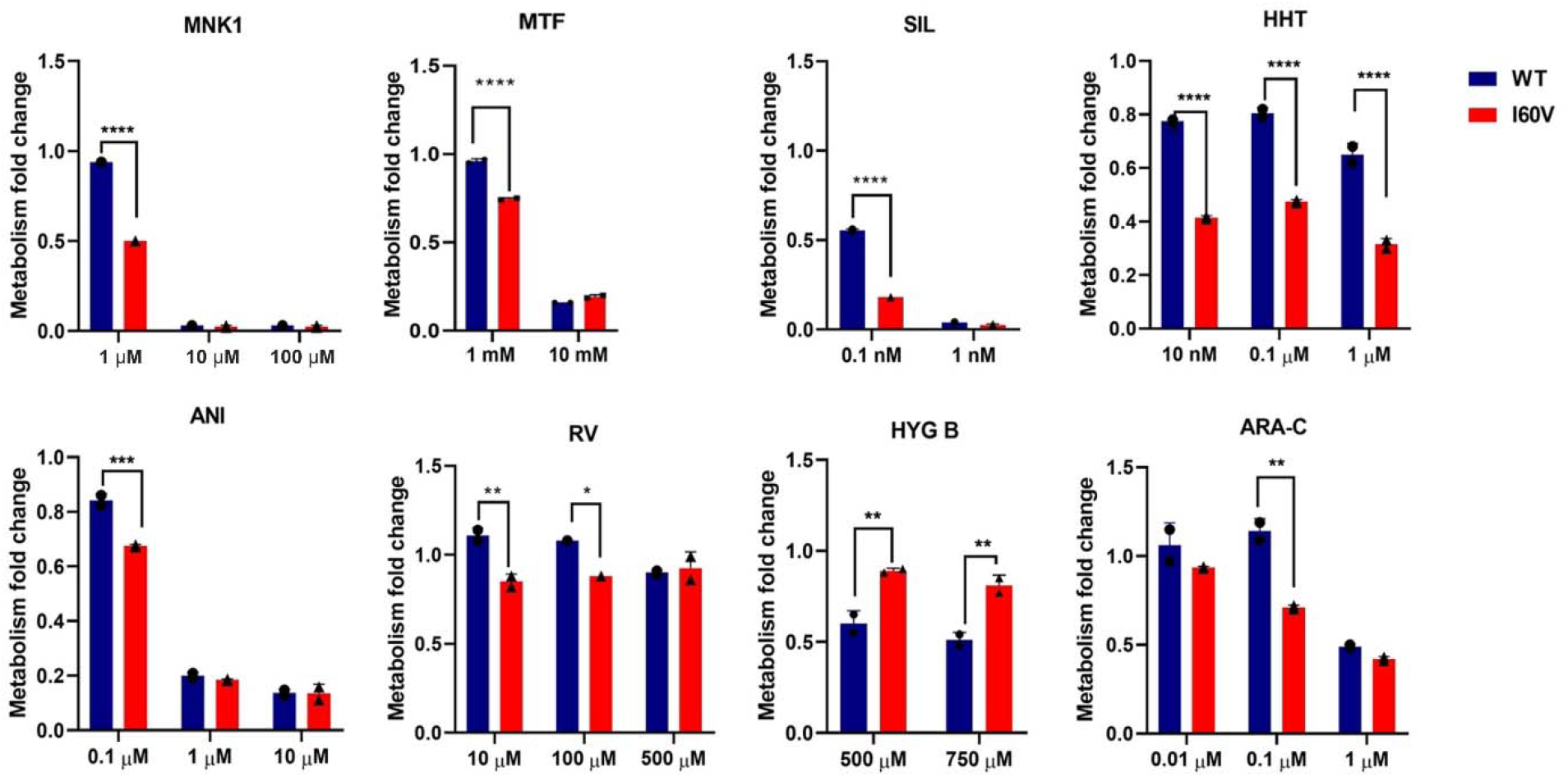
Impact of drug treatment on cell metabolic activity. The effect of the drugs on cellular metabolism was assessed by treating RPL5-WT (blue) and RPL5-I60V (red) clones with various anticancer drugs, primarily targeting translational machinery. Metabolic activity was evaluated, after a 48-hour treatment period, by AlamarBlue^™^ assay and data presented as fold change vs not treated sample. Graphs represent the average of two experiments run in duplicate. MNK1: MNK1 inhibitor; MTF: metformin; SIL: silvestrol; HHT: homoharringtonine; ANI: anisomycin; RV: resveratrol; HYG B: hygromycin B; ARA-C: cytarabine. Statistical significance was assessed by two-way ANOVA with Sidak’s multiple comparisons test *p<0.05, **p<0.01, ***p<0.001, ****p<0.0001.

#### 3.2.2 Effect of the mutation on drug-mediated modulation of protein synthesis

Because we tested various drugs known to inhibit ribosomal activity or protein synthesis, we aimed to evaluate how these drugs affect protein synthesis in cells with either wild-type RPL5 or the I60V mutation. To address this question, we performed non-radioactive labelling of newly synthesized proteins using puromycin incorporation (SUnSET) assay following a 24-hour drug treatment for both RPL5 WT and I60V clones.

Anisomycin and Homoharringtonine impacted on protein synthesis in both cellular types, with the mutant cells showing increased sensitivity compared to the WT cells (Fig. 4). Treatment with hygromycin B, another protein synthesis inhibitor, did not suppress protein production in either WT or mutant cells (Fig. 4). Instead, both cell types showed increased protein expression compared to untreated controls, suggesting resistance to the inhibitor. Notably, the mutant cells exhibited an even higher level of resistance than the WT. We then tested two inhibitors of translation initiation, a MNK1 inhibitor and Silvestrol. Both treatments determined an inhibition of protein synthesis compared to control conditions, albeit with no significant differences based on RPL5 genetic background. In addition, treatment with the antimetabolic agent cytarabine did not result in significant changes in protein synthesis levels in either the WT or I60V mutant clones (Fig. 4). Treatment with metformin, a compound targeting the mTOR signaling pathway, showed a significantly higher protein synthesis inhibition in the mutant cells compared to the WT cells, which in turn appear to be resistant to this compound (Fig. 4). Similarly, treatment with resveratrol determined no protein synthesis changes in WT cells, while I60V mutant cells showed a higher, albeit not significant, inhibition of protein synthesis (Fig. 4).

**Fig. 4.**
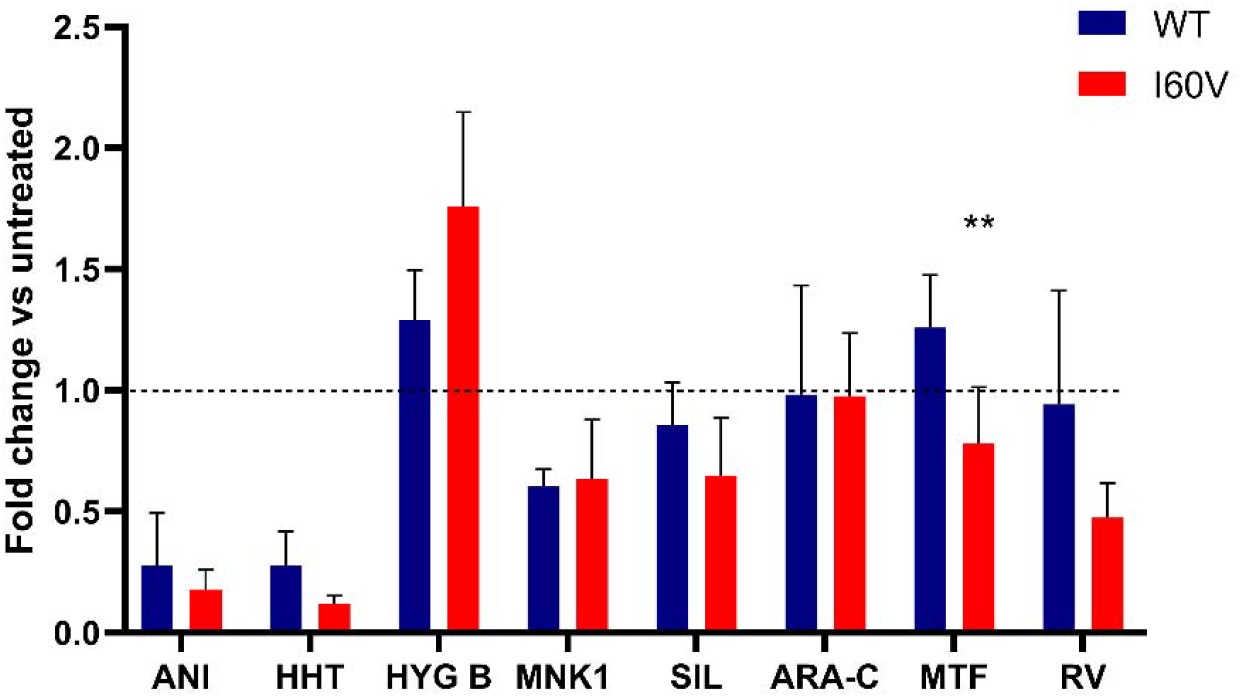
Effect of different compounds on protein synthesis. SUnSET analysis depicts the puromycin incorporation in cells treated for 24 hours with drugs. Anisomycin (ANI, 0.1µM); homoharringtonine (HHT, 10nM); hygromycin B (HYG B, 500µM); MNK1 inhibitor (MNK1, 1µM); silvestrol (SIL, 0.1nM); cytarabine (ARA-C, 0.1µM); metformin (MTF, 1mM) and resveratrol (RV, 10µM). The graph provides a quantitative representation of puromycin incorporation (representative western blot in Supplementary Figure 3), with results expressed as fold change relative to untreated cells + SD. Data represent the mean of two independent experiments. Statistical significance was assessed by t-test, **p<0.01.

#### 3.2.3 Effect of the mutation on drug-mediated modulation of cell viability, cytotoxicity and apoptosis

Given the known association between changes in metabolic activity and cell viability or apoptosis [47], further assays were conducted to validate what reported in the previous paragraphs, with the aim to evaluate the drugs’ efficacy in inhibiting proliferation in both RPL5-WT and RPL5-I60V clones. Optimal concentrations for viability assays were determined for each drug, based on their demonstrated efficacy in the metabolic assays (Fig. 3), and are reported in Figure 5A. Cells were treated for 72 hours, and, within this time frame, viability was assessed at various time points to ensure a comprehensive evaluation of the drugs’ effects. As shown in Fig. 5A, drugs impacted cell viability in both RPL5-WT and RPL5-I60V clones, but most drugs began to exhibit different and significant effects on clones only after 40 hours since treatment start. In general, it is possible to distinguish two different effects of the treatments on cells regardless of RPL5 genotype. In fact, while a reduction in viable cells number is observed after treatment with the MNK1 inhibitor, metformin, anisomycin, hygromycin B and cytarabine, a cytostatic effect is observed with the other drugs (silvestrol, homoharringtonine and resveratrol). MNK1 inhibitor and anisomycin showed the same trend (a high decrease after 16 hours and then a constant decrease in cell viability) on both wild type and mutant cells, with the latter showing higher sensitivity. In contrast, metformin, hygromycin B, and cytarabine demonstrated a comparatively moderate and temporally stable effect on cell viability. While metformin elicited minimal RPL5-dependent differences, hygromycin B and cytarabine induced more pronounced effects. Notably, I60V clones exhibited greater resistance to hygromycin B treatment compared to wild-type clones. Silvestrol and resveratrol exerted a predominantly cytostatic effect on mutant clones, whereas wild-type clones exhibited a modest increase in cell viability. Similarly, homoharringtonine exhibited a cytostatic effect on RPL5-I60V mutant clones, whereas wild-type clones displayed greater resistance to the compound. In summary, cell viability analyses revealed three distinct drug response profiles—sensitivity, resistance, and an intermediate cytostatic effect. Moreover, the RPL5 genotype appeared to influence the cellular response to treatment.

**Fig. 5.**
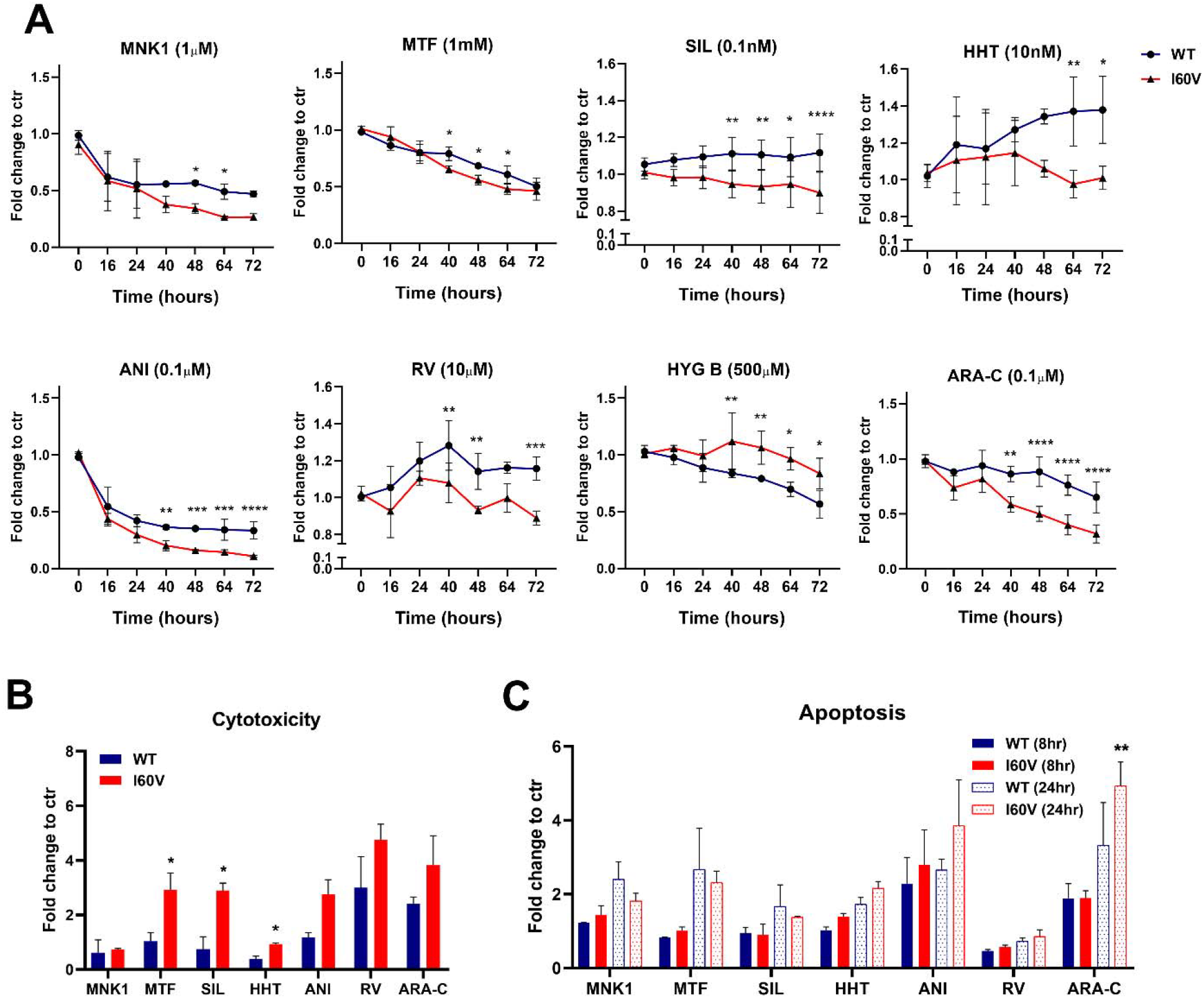
Impact on cell viability, cytotoxicity and apoptosis. (A) Cell viability was assessed with RPL5-WT (blue) and RPL5-I60V (red) clones using the RealTime-Glo^™^ assay (Promega). Cells were exposed to specified drug concentrations as indicated in figure. MNK1 inhibitor (MNK1, 1µM), Metformin (MTF, 1mM), silvestrol (SIL, 0.1nM), homoharringtonine (HHT, 10nM), anisomycin (ANI, 0.1µM), resveratrol (RV, 10µM), hygromycin B (HYG B, 500µM), and cytarabine (ARA-C, 0.1µM). Viability was monitored by measuring luminescence for 72 hours. Experiments were conducted in triplicate for each drug, with appropriate controls and data are represented as fold change vs control (ctr) ± SD. Statistical significance was assessed by two-way ANOVA with Sidak’s multiple comparisons test *p<0.05, **p<0.01, ***p<0.001, ****p<0.0001. (B) Cell cytotoxicity and (C) apoptosis were assessed in RPL5-WT (blue) and RPL5-I60V (red) clones using the ApoTox-Gloy^™^ assay (Promega). Cells were exposed to specified drug concentrations, consistent with those used for cell viability assays, for 40 hours to assess cytotoxicity and for 8 and 24 hours to assess apoptosis. Data represent the mean of two independent experiments run in duplicate and are expressed as fold change vs control (ctr) ± SD. Statistical significance was assessed by two-way ANOVA with Sidak’s multiple comparisons test *p<0.05, **p<0.01,***p<0.001, ****p<0.0001.

A comprehensive evaluation of treatment efficacy was conducted, considering the heightened sensitivity of RPL5-I60V cells to most tested compounds. However, due to the consistently diminished response of RPL5-I60V clones to hygromycin B across metabolic, viability, and protein synthesis assays, this compound was excluded from subsequent experiments. Utilizing the ApoTox-Glo^™^ assay, cytotoxicity and apoptosis were determined, thereby providing insights into cellular response. The results are summarized in figure 5B and C, focusing on the comparison between the responses of RPL5-WT versus RPL5-I60V treated cells. In our investigation of MNK inhibition, we observed induction of apoptosis without significant differences in cytotoxicity between wild-type (WT) and RPL5-I60V mutant cells. Metformin treatment resulted in a marked increase in cytotoxicity in I60V mutants, while apoptosis levels remained comparable across genotypes, suggesting activation of apoptosis independent of genotype-specific differences. Silvestrol had minimal impact on WT cells, showing no cytotoxic effect and only a slight increase in apoptosis at 24 hours. In contrast, I60V mutant cells exhibited reduced viability and elevated cytotoxicity following silvestrol treatment. Homoharringtonine elicited resistance in WT cells, whereas I60V mutants displayed a cytostatic response, accompanied by increased apoptosis and significantly elevated cytotoxicity compared to WT cells, despite no overall increase relative to untreated controls. Anisomycin exerted pronounced effects on both genotypes, with a stronger impact on I60V mutants, reflected in enhanced apoptosis and cytotoxicity. Resveratrol induced cytotoxicity in both WT and mutant cells, with a more substantial effect in I60V mutants, though apoptosis activation remained minimal in both. Finally, cytarabine treatment led to greater cytotoxicity in mutant cells, with apoptosis observed at both time points and significantly elevated in I60V mutants at 24 hours.

## 4. Discussion

First proposed over two decades ago [48], the concept of ribosomal heterogeneity as a regulator of protein synthesis has since gained substantial support in both physiological and pathological settings. Current evidence suggests that variations in ribosome composition—and consequently, structure—can drive functional specialization, enabling precise control of mRNA translation independently of transcription [23,49–52]. In pathology, ribosome diversity arising from aberrant RP sequence has been demonstrated to have functional repercussions on protein synthesis. It is, for instance, the case for RPL9 L20P in Diamond Blackfan Anemia, a ribosomopathy [26], or of RPL10 R98S in pediatric T-ALL [53]. In T-ALL, RPL10 mutations have been associated to altered translation, resulting in the rewiring of metabolic and stress response pathways [53–55]. In addition to the RPL10 R98S mutation—the most common and best characterized RP mutation driving disease in pediatric T-ALL—somatic mutations in RPL5 have also been reported, with missense variants being the most prevalent [56]. To investigate the role of RPL5 I60V missense mutation in ribosomal function within the context of T-ALL, we generated knock-in mutants in Jurkat cells, a pediatric T-ALL-derived cell line. RPL5 is a structural part of the central protuberance of the large subunit of the ribosome, that is involved in the functional interconnection between the decoding center in the 40S and the peptidyl transferase center in the 60S [57]. Thus, mutations in RPL5 can potentially alter ribosome translational activity. In addition to its structural role in the ribosome, RPL5 is critically involved in pre-rRNA processing and ribosome biogenesis, and sequence alterations may disrupt these functions as well [58]. We began our work by assessing the impact of the I60V mutation on ribosome biogenesis and function. Our analyses revealed that the mutation slightly impaired the biogenesis of the 60S ribosomal subunit. Mature ribosomes purified under high-stringency conditions from both wild-type (WT) and RPL5 mutant clones showed no compositional differences, apart from the I60V substitution. This included ribosomal protein content as well as site-specific rRNA modifications, specifically pseudouridylation and 2⍰-O-methylation, the two most abundant modifications in rRNAs [45]. Noteworthy, despite purifying ribosomes in highly stringent conditions, the MS analysis highlighted a differential enrichment in few non-ribosomal proteins, namely PTI-1, ARL4C, ERCC4, LRPPRC, NPC1, ABCB10, ETFDH, B3GNT2, and TRIM27 (Table 3). These proteins were present in much lower amounts than ribosomal proteins in the analyzed samples (several hundred times less abundant than ribosomal proteins by comparing iBAQ values). Among all these proteins, an effect on mRNA translation was previously reported only for PTI-1, whose expression has been associated to RPL5 and RPL4 [59]. Despite involving a single amino acid change, the functional impact of the I60V substitution within ribosomes was striking. Ribosomes incorporating I60V-RPL5 exhibited a marked increase in intrinsic protein synthesis activity, independent of the mode of translation initiation—whether 5⍰ cap-dependent or IRES-mediated. This enhanced translational capacity was accompanied by a faster proliferation rate in mutant cells, suggesting a growth advantage conferred by the mutation. To our knowledge, this is the first report of a RP mutant being incorporated into ribosomes, resulting in enhanced translational efficiency.

Based on these first results, we asked ourselves whether mutant cells could be differentially affected by treatment with compounds targeting ribosomes or, more in general, the translational machinery. Indeed, the localization of RPL5-I60V in the central protuberance of ribosomes could potentially influence sensitivity to several ribosome-targeting antibiotics, including homoharringtonine, hygromycin B, and anisomycin (all targeting the A site in the peptidyl transferase center). Leukemia cells heavily rely on cap-dependent translation via hyperactivation of signaling converging on eIF4A and eIF4E [60,61]. Since the RPL5-I60V mutation enhances translation rates, we hypothesized that these cells may exhibit heightened sensitivity to inhibitors of protein synthesis, like an inhibitor mitogen-activated protein kinase interacting kinase 1 (MNK1) [62], which targets phosphorylation of translation initiation factor 4E (eIF4E), and silvestrol, which targets eIF4A [60]. Moreover, we selected drugs targeting the mTOR signaling pathways, such as resveratrol [63] and metformin [64,65], which has also been shown to induce endoplasmic reticulum stress and inhibit the unfolded protein response (UPR) in leukemia cells [66]. For some of these compounds, prior use in experimental cancer therapies is reported (e.g. NCT01324180 for metformin; NCT00675350 for homoharringtonine, NCT00920803 for resveratrol, from www.clinicaltrials.gov). Additionally, we included cytarabine, a cornerstone chemotherapeutic agent in acute leukemia [67]. With the exception of cytarabine, all the tested compounds are expected to inhibit protein synthesis, albeit through distinct mechanisms as mentioned above. Indeed, our data show that only a subset of them— anisomycin, homoharringtonine, silvestrol, and the MNK1 inhibitor—was able to reduce protein synthesis to some extent in both genetic contexts. In contrast, metformin and resveratrol were effective only in the mutant background, suggesting that hyperactivated protein synthesis in RPL5-I60V cells may increase their reliance on this process. Furthermore, and in line with the metabolic data, mutant cells displayed even greater resistance to hygromycin B compared to WT.

When assessing the impact of the mutation on compound-mediated inhibition of cellular metabolism, we observed a bimodal response. Several drugs—such as silvestrol, anisomycin, MNK1 inhibitor, cytarabine, metformin, homoharringtonine, and resveratrol—exhibited greater efficacy in RPL5-I60V clones compared to RPL5-WT. In contrast, hygromycin B was less effective in RPL5-I60V cells. These findings suggest that the RPL5 mutation alone does not confer a general increase in resistance or sensitivity. Rather, drug-specific mechanisms of action, along with additional factors such as cellular uptake, metabolism, or clearance, likely contribute to the differential responses observed. In addition, while differential sensitivity to some compounds— such as homoharringtonine and hygromycin B—was evident across the entire tested concentration range, for most drugs it was restricted to the lower concentrations. This suggests that the mutation modulates sensitivity rather than producing an all-or-none effect.

Most drugs exhibited a significant impact on cell viability, in line with findings from the metabolic assays. While some compounds—such as silvestrol, homoharringtonine, and resveratrol—exerted a cytostatic effect, others—including the MNK1 inhibitor, metformin, anisomycin, hygromycin, and cytarabine—led to a marked reduction in cell number as early as 16 hours after treatment initiation, suggesting the activation of cell death pathways. It worth underscoring, however, that the effect on cellular viability was, in most cases, genotype-dependent, with the I60V mutants being generally more sensitive compared to WT RPL5 genotypes. Consistently with the findings on cellular metabolism and protein synthesis inhibition, the only exception was represented by hygromycin B, for which mutant cells appeared to be more resistant compared to the WT genotype. Dependent of the treatment, we observed the activation of different cell death pathways. Most compounds, except for the MNK1 inhibitor and homoharringtonine, showed a cytotoxic/necrotic effect at the second day of treatment, with mutant cells being more prone to the activation of this pathway. On the other hand, treatments also tuned up the apoptotic pathway, with a clear effect after 16 hours except for silvestrol and resveratrol. Mutant cells appeared to activate apoptosis to equal or higher levels compared to RPL5-WT cells. Our data are in line with previous literature showing a proapoptotic effect on leukemia cells of the different compounds in the tested concentration ranges ([68] ARA-C; [69] ANI; [70] HHT; [71] MTF; [72] MNK1).

Although the data presented here suggest a ribosome-centered effect of the I60V mutation, we cannot exclude the possibility that this mutation also affects ribosome-independent functions of RPL5. These may include its role as an activator of ribosomal stress responses via the 5S-ribonucleoparticles (5S-RNPs) [73] as well as its involvement in transcriptional regulation [15,74]. Future studies will be needed to address this question and to investigate the potential of combination therapies using the tested compounds with distinct mechanisms of action, whose efficacy appears enhanced in the context of RPL5-I60V.

## 5. Conclusions

This study opens a new perspective in the treatment of those cancers which bear alterations in ribosomes or in the translational machinery, shedding light on the potential of ribosomal proteins as therapeutic targets for the development of increasingly personalized, more effective and less toxic therapies.

## 6. Authors contributions

Conceptualization, funding acquisition, supervision: MP; data curation, investigation, formal analysis: DP, IJR, MD, AMH, YC, YM, VM, KF; Visualization: LB, MP, DP, IJR, MD, AMH, YC, YM, VM, KF; Writing: LB, IJR, YC, YM, MP.

## 7. Acknowledgments

This article/publication is based upon work from COST Action TRANSLACORE CA21154, supported by COST (European Cooperation in Science and Technology). MP warmly thanks Professor Lorenzo Montanaro and his research group for continuous discussion on the project, and Prof. Kenneth B. Marcu for critical reading of the final draft of the manuscript.

